# Flexibility of a mammalian lipidome — insights from mouse lipidomics

**DOI:** 10.1101/2021.05.12.443735

**Authors:** Michał A. Surma, Mathias J. Gerl, Ronny Herzog, Jussi Helppi, Kai Simons, Christian Klose

**Affiliations:** Lipotype GmbH, Tatzberg 47, 01307 Dresden, Germany; MPI-CBG, Pfotenhauerstraße 108, 01307 Dresden, Germany

**Keywords:** lipid, lipidomics, mass spectrometry, mouse, murine, *Mus musculus*

## Abstract

Lipidomics has become an indispensable method for the quantitative assessment of lipid metabolism in basic, clinical, and pharmaceutical research. It allows for the generation of information-dense datasets in a large variety of experimental setups and model organisms. Previous studies, mostly conducted in mice (*Mus musculus*), have shown a remarkable specificity of the lipid compositions of different cell types, tissues, and organs. However, a systematic analysis of the overall variation of the mouse lipidome is lacking. To fill this gap, in the present study, the effect of diet, sex, and genotype on the lipidomes of mouse tissues, organs, and bodily fluids has been investigated. Baseline quantitative lipidomes consisting of 796 individual lipid molecules belonging to 24 lipid classes are provided for 10 different sample types. Furthermore, the susceptibility of lipidomes to the tested parameters is assessed, providing insights into the organ-specific lipidomic plasticity and flexibility. This dataset provides a valuable resource for basic and pharmaceutical researchers working with murine models and complements existing proteomic and transcriptomic datasets. It will inform experimental design and facilitate interpretation of lipidomic datasets.

## 1 Introduction

Lipidomics is the quantitative and comprehensive analysis of lipids in biological samples [Shevchenko and Simons, 2010, Wang et al., 2016]. By means of automated sample extraction, state-of-the-art mass spectrometry, sophisticated and validated spectra annotation and data analysis processes as well as modern statistical methods (machine learning), biologically relevant information can readily and reliably be obtained based on thousands of data points O’Donnell et al. [2020]. Therefore, lipidomics has become an indispensable tool for understanding (lipid) metabolism at the molecular level in basic, clinical, and pharmaceutical research. In the clinical context, the analysis of human blood samples in large population studies have revealed novel lipid signatures for a variety of indications such as: diabetes type 2 [Fernandez et al., 2019, Rhee et al., 2011], obesity [Beyene et al., 2020, Gerl et al., 2019], cardiovascular diseases [Tabassum et al., 2019, Rämö et al., 2019, Laaksonen et al., 2016, Ottosson et al., 2021] and neurological disorders [Penkert et al., 2020]. Furthermore, in nutritional research lipidomics has proven a powerful read-out in dietary intervention studies [Kessler et al., 2020].

Lipidomics has successfully been applied to study disease and disease-related mechanisms in many different indications. Among these are neurological disorders [Bosch-Queralt et al., 2021, Cascalho et al., 2020], liver disease [Parker et al., 2019] and cancer [Bi et al., 2019, Peck et al., 2016, Saliakoura et al., 2020], resulting in the identification of potential drug targets involved in lipid metabolism [Matsushita et al., 2021].

A primary model for these studies is the mouse *Mus musculus*. Typically, lipidomic changes in blood plasma and a variety of tissues or organs are assessed to understand disease mechanisms or modes of drug action. Several studies have shown that different cells, tissues, and organs exhibit highly specific lipid composition [Fitzner et al., 2020, Jain et al., 2014, Parker et al., 2019, Pradas et al., 2018, Symons et al., 2021]. However, the systematic knowledge of the natural, biological lipidomic variation of different organs and bodily fluids is lacking, hindering interpretation of such studies.

To fill this gap, in the present study, the effect of diet, sex, and genotype on the lipidomes of mouse tissues, organs, and bodily fluids has been investigated. Baseline quantitative lipidomes consisting of more than 796 individual lipid molecules for 10 different sample types (full blood, blood plasma, liver, skeletal muscle, brain, kidney, adipose tissue, small intestine, lung, and spleen) are provided. Furthermore, susceptibility of lipid levels to the tested parameters is assessed, providing insights into the organ-specific lipidomic plasticity and flexibility of a mammalian organism. This dataset provides a valuable resource for basic, pharmaceutical, and clinical researchers using mouse as model system and complements existing proteomic and transcriptomic datasets. It will inform experimental design and facilitate interpretation of lipidomic datasets.

## 2 Results

### 2.1 Study objective and design

The aim of the present study is to investigate the lipidomes of mouse tissues, organs, and bodily fluids and how they are affected by different diets in mice of different genotype and sex. To this end, two standard laboratory mouse strains were selected, representing genetically outbred and inbred populations. Parental mice were allowed to breed, and females (already during pregnancy) and the consecutive litter mice were fed with a high protein (18 weight%) or a low protein (14 weight%) diet; both diets representing healthy and standard compositions. For each combination of conditions, three mice of each sex were sacrificed, and 10 sample types collected, prepared for lipid extraction, and analysed using a quantitative, high-throughput shotgun lipidomics platform. Sample types included: full blood, blood plasma, liver, skeletal muscle, brain, kidney, adipose tissue, small intestine, lung, and spleen; in total 240 samples. For details see Materials & Methods.

The present study design enabled a factorial analysis of the influence of the different conditions (i.e., genotype, diet, and sex) on each individual lipid in every sample type. The comprehensive set of sample types allowed for a detailed description of mouse organ lipidomes regarding abundances of lipid classes and individual lipid molecules, fatty acid saturation/unsaturation, chain length and hydroxylation profiles as well as their susceptibility to differences in the experimental conditions. Furthermore, changes of lipids in different organs or tissues across the different conditions could be correlated with changes in blood lipidomes (full blood or blood plasma). This allowed to investigate, whether blood lipidomes are useful proxies for various organ and tissue lipidomes.

### 2.2 Method validation and analytical performance

We have published detailed methods for mass spectrometry-based shotgun lipidomics of blood plasma [Surma et al., 2015] and adipose tissue [Grzybek et al., 2019]. We have shown that for these two matrices the analytical precision (repeatability) is well below 15% (relative standard deviation, RSD), sensitivity is in the nM range and the linear dynamic range spans four orders of magnitude.

To further extend method validation for the present sample types, liver, brain, and full blood were chosen as representative material to assess repeatability, sensitivity, and dynamic range of the analytical method. These particular sample types are the most complex and exemplary regarding lipid composition and quantities to be encountered across various tissues, organs, and other samples. As expected for a shotgun lipidomics method, sensitivity was in the nM range and linearity was achieved over four orders of magnitude. Repeatability as assessed by the extraction and analysis of aliquots of a representative sample, was 5.6%, 6.3%, and 8.3% RSD for liver, brain, and full blood samples, respectively. For details see Supplemental Material.

The sample set was analyzed in four analytical batches, according to the required extraction and mass spectrometry methods (see Materials & Methods). The analytical performance was assessed by including sample type-specific reference samples. The median RSD on the level of individual lipid (sub)species for plasma was 6.6%, for full blood 15.7%, for adipose tissue 4.9%, and for the remaining tissues 12.3%. A total of 24 lipid classes comprising a total of 796 lipid species could be quantified across all sample types (Table 1). Among these, spleen and lung exhibited the broadest coverage (24 lipid classes), while in adipose tissue the lowest number (11) of lipid classes could be detected. The reproducibility as assessed by the median RSD for lipid (sub)species of the biological triplicates of each organ and combination of experimental conditions was between 12.5% and 25.8%. Reproducibility was highest for brain and full blood and lowest for kidney and spleen. Note, that these reproducibility values include both technical variation of the analytical method and biological variation between the triplicate mice.

**Table 1:**
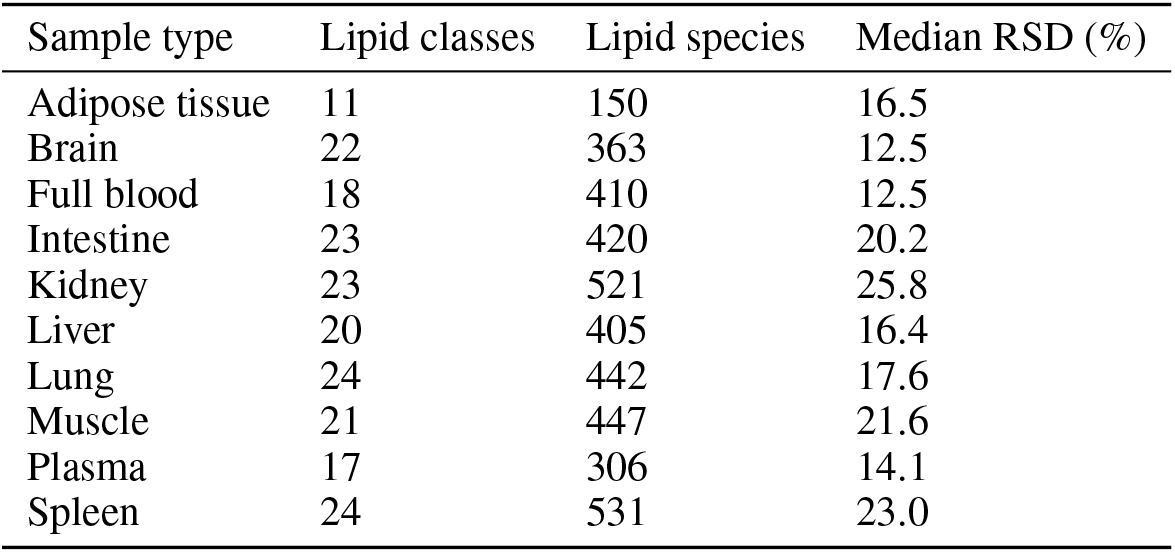
Lipidome coverage and reproducibility of the measurements. Median RSD is provided for individual lipid (sub)species in the biological triplicates.

### 2.3 Lipidomic characteristics of mouse organs and tissues

To obtain an overview of the similarities between different sample types, a principal component analysis (PCA) was performed based on amounts of lipid species normalized to total lipid for all experimental conditions (molar fractions expressed as mol%, Figure 1 A). Unsurprisingly, almost every sample type forms a separate cluster, indicating a distinct, organ-specific lipid composition. Notable exceptions are adipose tissue and muscle, which are not discriminated well in the first two principal components (PC) of the PCA. This lipidomic similarity of adipose and muscle tissues, despite their apparent biological difference, is caused by the fact that both samples contained high (*>* 90 mol%) amounts of TAG^1^. As this observation is obvious for adipose tissue, for the muscle it is most likely caused by not differentiating between muscle tissue and the intermuscular fat during sample collection. Brain appears to have the most distinct lipid composition of all organs and tissues analyzed, as its samples are clustered well separated from other sample types, in both principal component dimensions (PC1 and PC2). Furthermore, along PC1, brain and adipose tissue/muscle are most distant to each other, indicating distinct compositional differences due to the high concentrations of TAG in muscle and adipose tissue, which on the other hand is almost entirely absent in brain. Instead, brain exhibits high concentrations of cholesterol, PS, and most importantly, PE O- and hexosyl ceramide (Figure 1 D). Intestine samples form a distinct cluster separated from the other sample types in PC2. They show high concentrations of PC O-, LPC, ceramides, LPE, and other lyso-lipid classes. With around 5 mol%, the LPE concentration in the intestine is by far the highest among all tissues and organs. Lung samples form another well-separated cluster. They are rich in sterols, SM and in particular PG. Like lung, spleen is rich in cholesterol (ca. 25 mol%, Figure 1 D). In addition, in spleen there are significant amounts of PE O-, PC O-, PI, PS, and SM. The blood samples (full blood and plasma) obviously share characteristic similarities, such as high concentrations of CE and LPC. Full blood differs from plasma because it contains higher concentrations of PE, PE O-, PI, and PS, which are contributed by the cellular components of blood, mainly erythrocytes. Interestingly, kidney and liver samples are forming partially overlapping clusters. Liver and kidney, both share a comparably high concentration of DAG and intermediate concentrations of TAG and cholesterol. Furthermore, liver has the highest PI concentration among all sample types.

**Figure 1:**
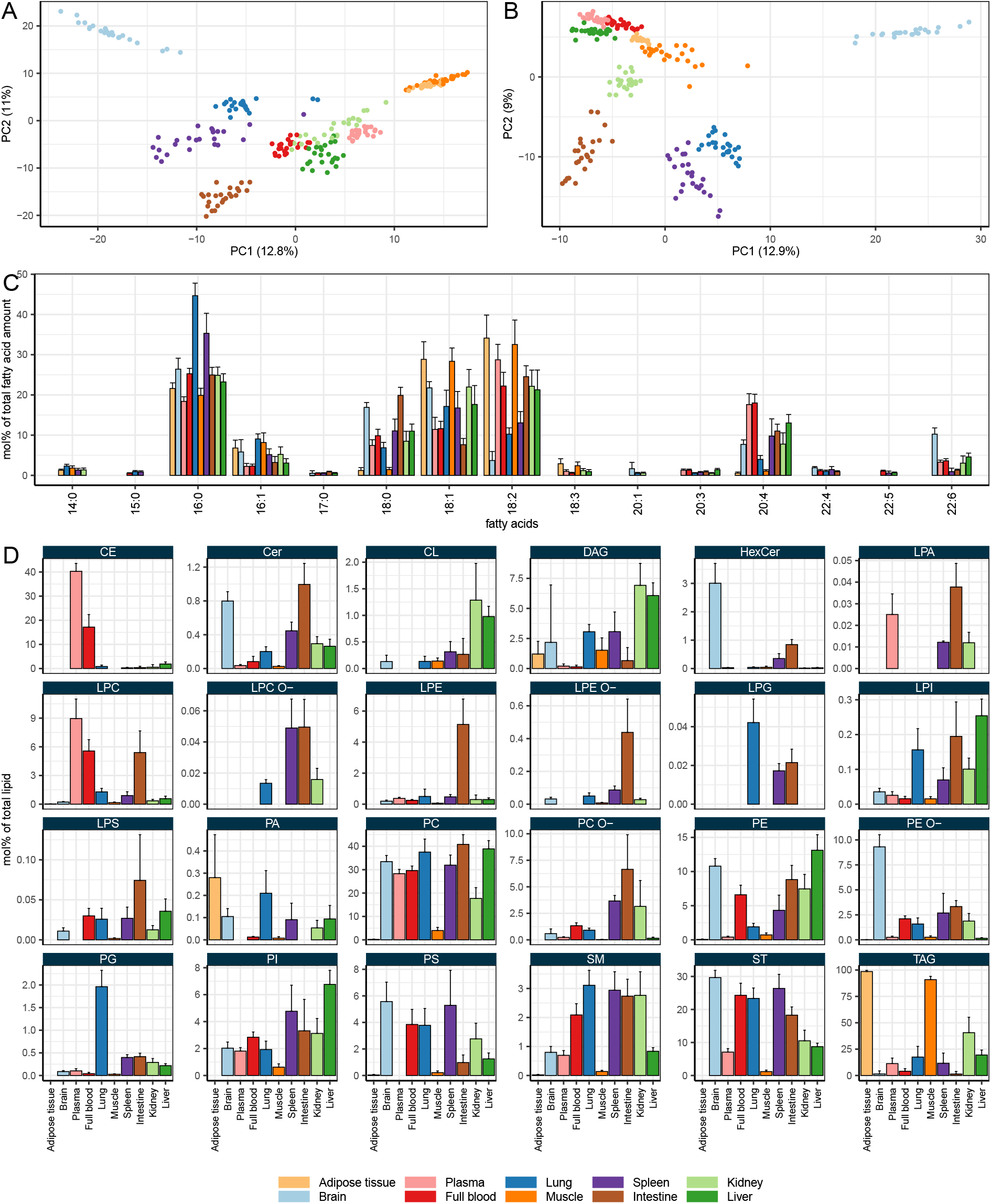
Lipid composition of mouse organs. A: Principal component analysis (PCA) based on lipid species in mol%. Axis labels indicate principal component (PC) 1 and 2, including % variance explained. B: PCA as in A, but excluding storage lipids (DAG, TAG, CE). C: Fatty acid profiles. D: Lipid class composition. Shown are mean values for all combinations of experimental conditions (biological triplicates for all combinations of diet, sex, and genotype; *n* = 24). Error bars indicate standard deviations.

When the storage lipids (CE, DAG, TAG) are excluded from the PCA, the tissue-specific clusters are in most cases preserved (Figure 1 B). This is the case for brain, kidney, intestine, spleen, and lung. For the remaining tissues, the clusters become even more distinct arguing for a tissue-specific composition of the membrane lipids in particular. This effect is most pronounced for adipose and muscle tissue which overlap entirely in the presence of storage lipids but form separate clusters based on membrane lipids.

The tissue-specific lipid (sub-)species composition is based in part on a distinct fatty acid profile of the different sample types (Figure 1 C). In general, the three most abundant fatty acids are 16:0 (palmitic acid), 18:1 (likely oleic acid), and 18:2 (likely linoleic acid). Palmitic acid is most abundant in lung tissue, where it is a major component of surfactant lipids. Plasma and full blood samples contain the highest amounts of the poly-unsaturated fatty acid (PUFA) arachidonic acid (20:4) while brain is rich in DHA (22:6). Brain also contains, together with intestine, comparably high amounts of stearic acid (18:0). Complete lipidomic data are provided in Supplemental Material.

### 2.4 Qualitative lipid class composition

Lipidome data can be presented in many ways. This is due to the fact, that lipids are built from structural entities that are shared among all lipid classes. This allows for grouping the lipidomic data according to different structural features. These structural features can be headgroups, the defining characteristic of lipid classes, or fatty acids with varying length (number of carbon atoms in an acyl moiety) and number of double bonds (unsaturation). Therefore, the lipid class abundance (as the molar fraction of the total measured lipidome, expressed in mol%), the weighted mean of the number of double bonds, and the weighted mean of the number of carbon atoms in the fatty acid moiety of the lipids belonging to a given lipid class are three lipidomic features that enable a quantitative as well as qualitative description of a given lipid class. This results in a reduced number of lipidomic features. For example, the most complex sample type in the present study (spleen) contains 531 lipid molecules belonging to 24 lipid classes. By describing a lipid class by the three parameters, the number of features (the data granularity) to be analyzed is reduced to 72 (24 lipid classes *×*3), a seven-fold decrease. The advantage of condensing the lipidome data in this way is, that the number of features to be analyzed statistically can be decreased without losing the most important biological information, conveyed by lipid classes content and their length/saturation indices.

This condensed qualitative view on the lipid class composition can be displayed by plotting the weighted mean of the number of double bonds and the weighted mean of the number of carbon atoms in the fatty acid moiety of the lipids belonging to a given lipid class against each other (Figure 2).

**Figure 2:**
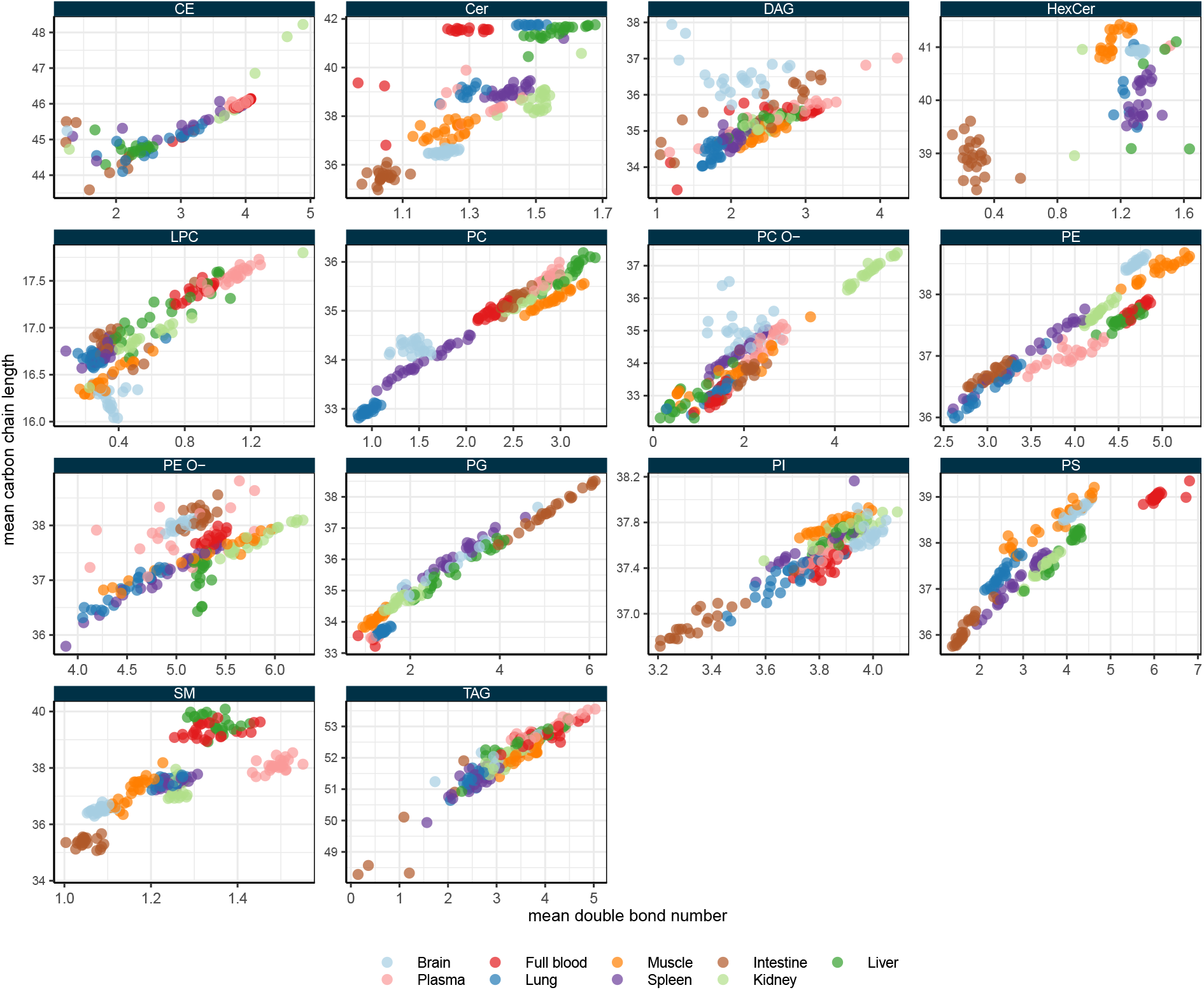
Feature profiles of the major lipid classes. Shown is the weighted mean number of double bonds per lipid class in a given sample and the weighted mean number of carbons in the hydrocarbon chain moiety per lipid class in a given sample (each dots represents an individual sample). Minor lipid classes (lyso-lipids) are omitted for clarity.

Unsurprisingly, for most lipid classes, there is a correlation between the carbon chain length and the degree of unsaturation: the longer the fatty acids, the more double bonds are to be found. However, this phenomenon is more pronounced in some classes than in others. For PC and PG there seems to be a perfectly linear relationship between chain length and unsaturation. However, the sphingolipids (ceramide, SM and hexosyl ceramide) do not follow this correlation but exhibit rather organ-specific length and unsaturation profiles due to the differential expression of ceramide synthases. For example, SM has a very distinct profile in intestine samples with short (35-36 carbons) and saturated (1-1.1 double bond) SM species. In plasma however, SM is longer (ca. 38 carbons) and more unsaturated (ca. 1.5 double bond).

A striking feature of the lipid class length and unsaturation profiles is that they appear to be more conserved in certain organs than in others. For example, especially brain exhibits a very distinct lipid class profile for the sphingolipids, PC, PE, PE O-, PI and PS. Similarly, lung shows a very tightly controlled lipid profile in PC and PG. Of note, lung PC exhibits the shortest and most saturated profile across all sample types analyzed. Intestine is another organ with rather specific lipid class profiles: its PI and PS are comparably short and saturated, while PE O-in intestine samples shows longer carbon chains (ca. 38) than in any other sample type.

Sphingolipids like ceramide, SM, and hexosyl ceramide show a specific chain length and double bond profile in basically every organ. Other lipid classes have a more distinct lipid class composition in some organs but can vary broadly in terms of fatty acid chain length and degree of unsaturation in others. A noteworthy example is PC, which in brain and lung has a very specific lipid profile, while especially in spleen, but also in plasma, its lipid class composition is rather variable. Similar observations can be made for other lipid classes, such as PE, PE O-, and PS.

One may generalize these observations and say that across all organs and tested conditions certain lipid classes exhibit a greater heterogeneity in their fatty acid composition (profile) than others. For example, PS shows a very broad double bond range (from 0 to almost 7) while PI shows a very narrow double bond range (from 3.2 to 4.1). Similarly, PC O-has a very broad carbon length range (from 32 to almost 38) while PE shows a very narrow carbon length range (from 36 to 39).

### 2.5 Plasticity of lipid classes

To express these observations in a simple parameter, we propose the concept of “lipidomic plasticity”. The lipidomic plasticity expresses the capability of a species (such as mouse) to adopt the amount and/or composition of lipidomic entities (such as lipid classes) in response to internal and external stimuli. For example, a lipid class that is observed to assume a very broad range of double bond numbers, carbon chain length, or abundances across a variety of experimental conditions, has a high degree of lipidomic plasticity. On the contrary, a lipid class which exhibits hardly any variation in abundance, double bond profile or carbon chain length has a low degree of lipidomic plasticity. Hence, plasticity is the degree of structural heterogeneity/variation within a given lipid class.

The plasticity of a lipid class was estimated as the product of the range of the weighted mean of the number of double bonds and the weighted mean of the number of carbon atoms in the hydrocarbon moiety of a lipid class (Figure 3 A, for details of the calculation see Materials & Methods). As defined above, a lipid class that exhibits a wide range of unsaturation and/or carbon chain length will have a high plasticity, while a lipid class with low variability will have a low plasticity. In this way, it is possible to easily compare the capability for adjusting the species profile of a lipid class across the different organs. For example, PC has a high plasticity in spleen, while very low in intestine (Figure 3 A). This is explained by the fact that PC clusters tightly around 35-35.5 carbons in the fatty acid moiety and 2.5 double bonds in intestine, while in spleen the ranges are from ca. 33 to 34.5 for carbons in fatty acids and double bond numbers between 1 and 2 (Figure 3 A). The analysis of lipid class plasticity therefore reveals, that in some organs the fatty acid composition of certain classes is more constrained (i.e., less affected by diet, sex, genotype) than in others (Figure 3 B). In brain, but also in lung, the lipid class profiles are the least plastic. Other organs, like spleen, tend to have more plastic lipid classes profiles. Liver shows a similar tendency for many lipid classes, but not for all.

**Figure 3:**
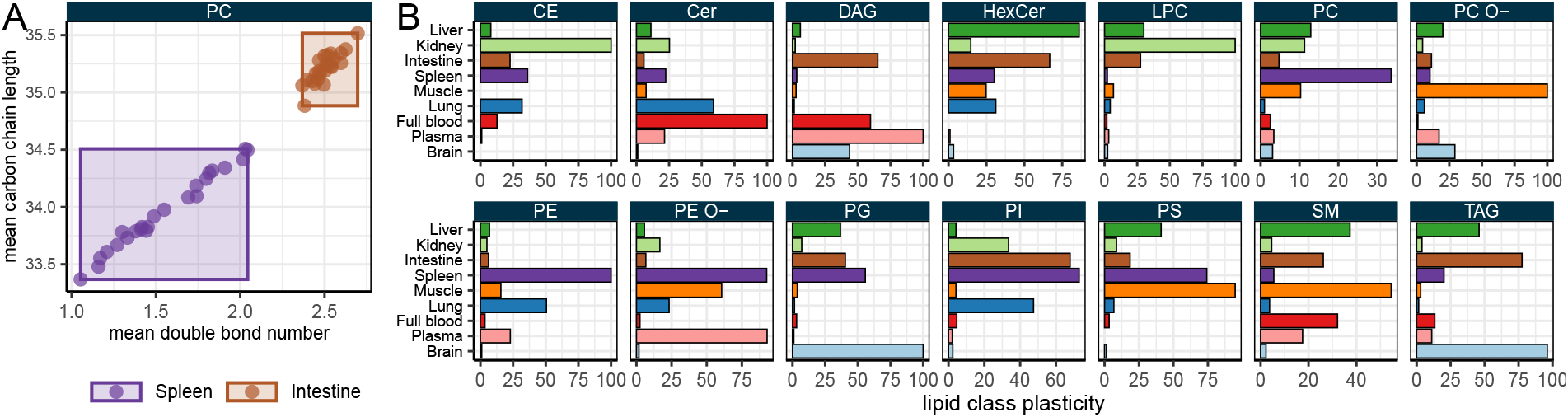
Lipid class plasticity across all combinations of experimental conditions. A: Plasticity is calculated per class and organ by multiplying the scaled ranges of values of weighted mean number of double bonds and the weighted mean number of carbons in the hydrocarbon chain moiety per lipid class. The wider the ranges, the higher the plasticity. B: Lipid class plasticity for the different mouse organs. For details see Materials & Methods.

In summary, along the parameters of lipid class abundance, carbon chain length and double bond profile, each lipid class exhibits a distinct quantitative and qualitative pattern in every organ. Here we introduce plasticity as an additional defining characteristic of each lipid class.

### 2.6 Flexibility of the mouse lipidome

As shown above, there is significant variation of the organ lipidomes across the tested conditions. An analysis of variance based on multiple linear regression with lipid species concentrations in mol% as outcome and sex, diet, genotype, and sample type as covariates shows that the major source of variance in the dataset is the sample type, accounting for about 60% of overall variance (not shown). This confirms the observations in the PCA (Figure 1). Therefore, organ identity is the most determining factor for lipid composition. Diet, sex, and genotype contribute to only about 5% of overall variance each, indicating that, in the range tested, their influence is more subtle.

To investigate effects of diet, sex, and genotype on the mouse organ lipidomes, multiple linear regression with lipidomic features (i.e., lipid class concentration in mol%, weighted mean of lipid class unsaturation, weighted mean of lipid class chain length) as outcome was performed. Out of 68 lipidomic features included in the analysis, 56 (82%) were significantly affected by any of the tested conditions in at least one of the organs. Those lipidomic features that were not affected represent only minor, less abundant, lipid classes, such as lyso-lipids. More specifically, every organ is affected by diet, sex, and genotype in at least one lipidomic feature. The only exception is brain, which is not affected by diet and sex. However, genotype has a highly specific effect on the brain sphingolipids and cholesterol (Figure 4 B). Cholesterol, SM, and hexosyl ceramide content is increased in outbred mice, while ceramide content is decreased. Moreover, unsaturation and carbon chain length are reduced for ceramides and SM in outbred mice, while the degree of unsaturation of hexosyl ceramide is increased. Complete results for the regression analysis are provided in Supplemental Material.

**Figure 4:**
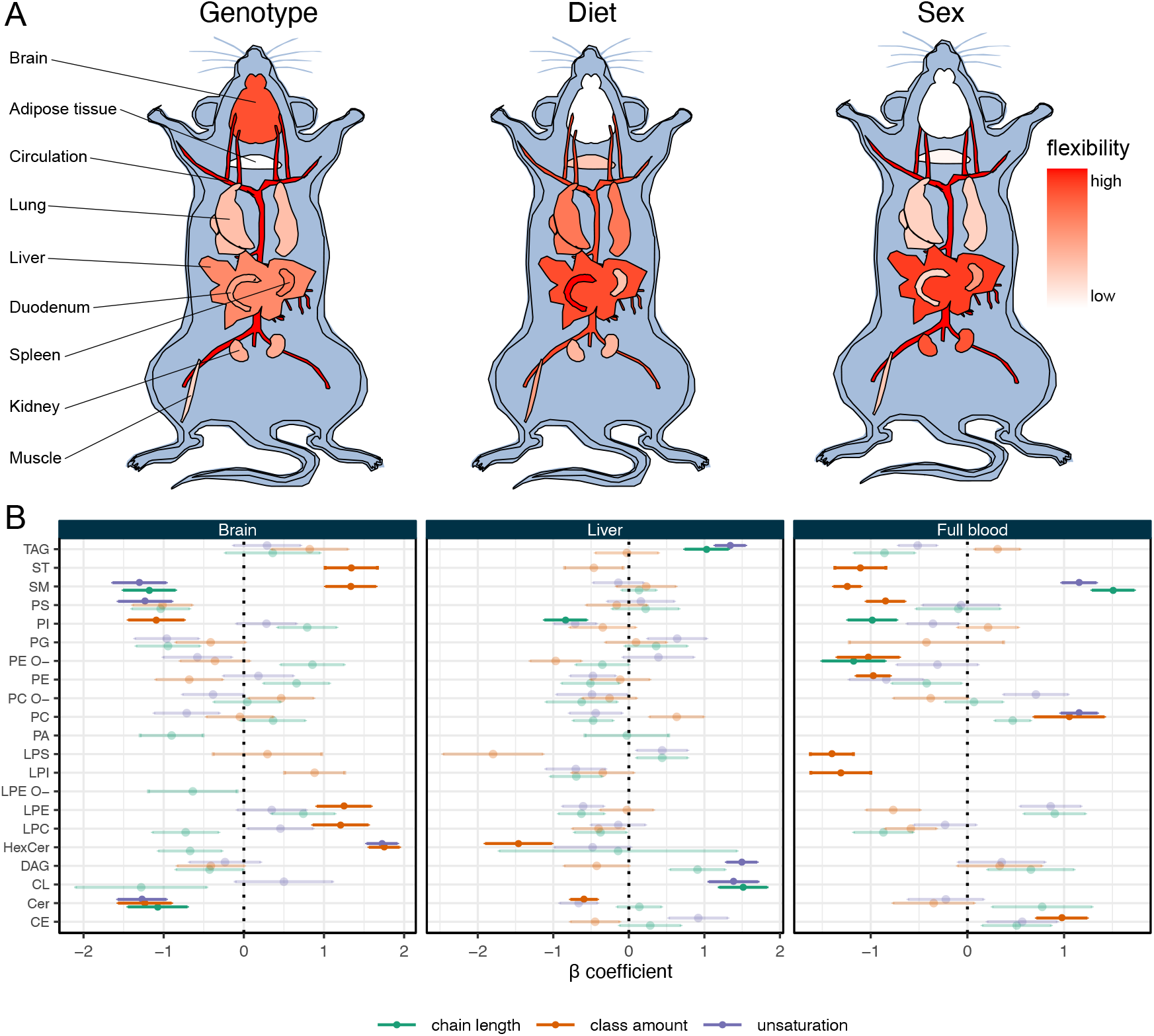
Flexibility of the mouse lipidome. A: Analysis of lipidomic flexibility in mouse organs based on the tested experimental conditions. For details see main text. B: Analysis of lipidomic responses by multiple linear regression. Shown are *β* coefficients. Error bars indicate standard error. Examples shown are: Genotype effects for brain; diet effects for liver; sex effects for full blood. Significantly affected features (*p <* 0.01) are displayed non-transparently.

To summarize the lipidomic changes induced by the experimental conditions and to quantitatively assess their impact on the mouse organ lipidomes we apply the concept of lipidomic flexibility [Klose et al., 2012]. Lipidomic flexibility is defined as the magnitude of lipidomic changes induced by certain experimental conditions. Here, it is calculated by summing the absolute effect sizes (*β* coefficients of the linear regression) for significantly affected features (*p <* 0.01) of the different organs (Figure 4 A). In this way, the highly specific effect of the genotype on the brain lipidome becomes obvious. Furthermore, genotype and sex exert organ-specific effects. While plasma, full blood and brain are strongly affected by genotype, adipose tissue and lung are only weakly affected. Similarly, sex strongly influences the full blood lipidome (Figure 4 B), while brain, adipose tissue, but also lung and intestine are hardly affected. Also, the kidney and liver lipidomes exhibit sex-specific differences. In liver, diet has a significant impact on TAG and CL composition (Figure 4 B). Of note, full blood and liver usually show the highest flexibility, irrespective of the tested condition. Full blood (and in a similar way blood plasma) is strongly affected throughout the entire lipidome. There are significant changes not only in the storage lipid TAG and CE, but also in the major phospholipid classes PC, PI, PS as well as in the sphingolipid SM and cholesterol (see Supplemental Table). Hence, full blood can serve as a rich source of information on systemic changes in lipid metabolism.

### 2.7 Genotype- and sex-specific dietary effects on the lipidome

In this study, mice of each genotype and sex were fed two different diets: high (18%) and low (14%) protein (for details see Materials & Methods). This factorial design allows for the analysis of differential effects of a given condition depending on the status of the other conditions, in particular diet-induced lipidomic changes in dependence of genotype or sex. For example, we observed that ceramide levels in kidney samples are increased in inbred mice (female and male) fed a high protein diet as compared with low protein diet while they are not affected in kidneys of outbred mice (Figure 5 C). Similarly, the degree of unsaturation of PE is negatively affected by high protein diet in inbred mice but not significantly affected outbred mice.

**Figure 5:**
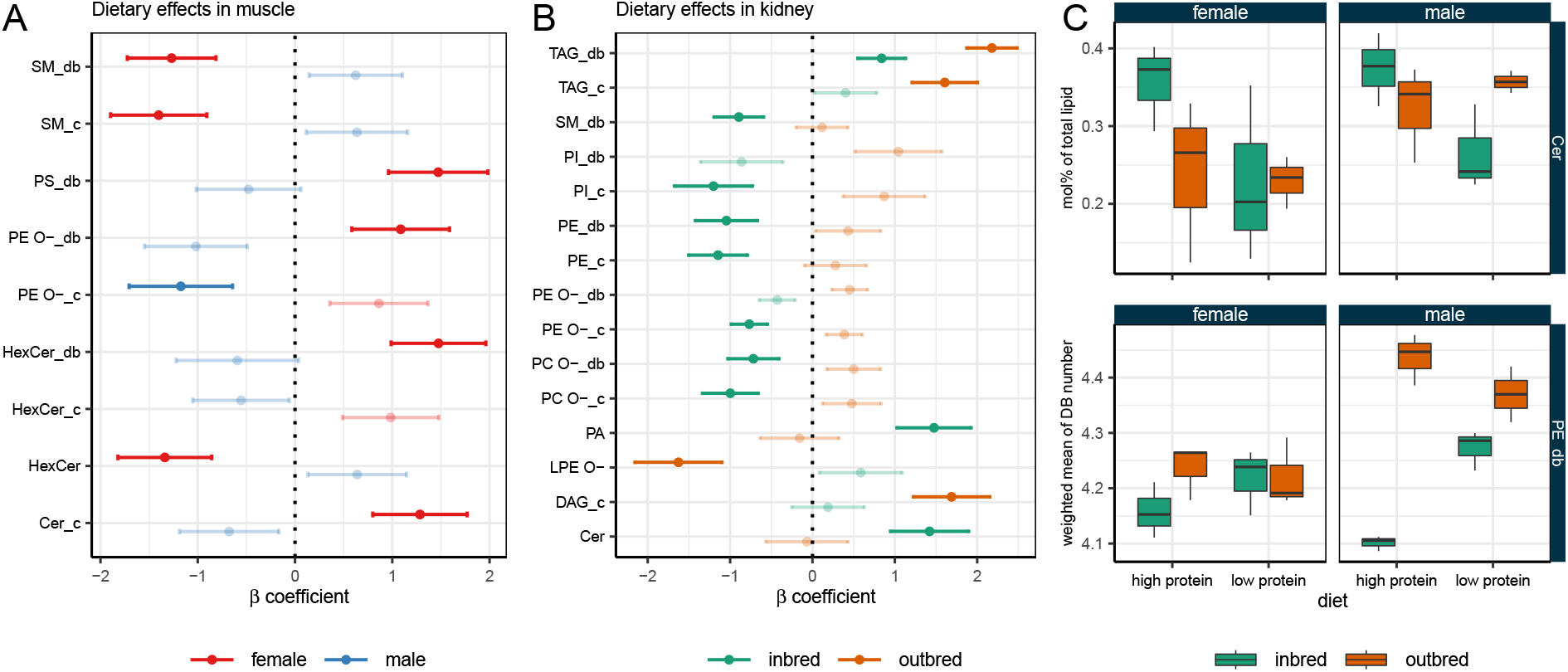
Sex-(A) and genotype-specific (B) effects of diet based on a multiple linear regression including interaction between diet and genotype and diet and sex, respectively. Shown are *β* coefficients. Error bars indicate standard error. Significantly affected features (*p <* 0.05 for the interaction term) are displayed non-transparently. Lipidomic features are depicted on the y axis (with “_db” denoting the weighted mean double bond number and “_c” denoting the weighted mean of the carbon chain length of a lipid class; when neither is specified, the feature name refers to lipid class abundance). C: Genotype-specific effects of diet on ceramide levels and degree of PE unsaturation in kidney.

We therefore performed a systematic analysis of sex-or genotype-specific effects of diet based on interaction terms in a multiple linear regression model. For example, sex-specific diet effects could be mostly detected in intestine and muscle. Of note, in muscle, differential effects could mostly be observed for the sphingolipids SM, hexosyl ceramide, and ceramide (Figure 5 A). Genotype-specific diet effects could be detected mostly in brain, intestine and kidney. In kidney, differential effects can be observed throughout many lipid classes, in particular phospholipids (Figure 5 B). Especially the degree of unsaturation and hydrocarbon chain length of ether lipids (PC O- and PE O-) change in opposite directions depending on the genotype. Similar effects could be observed in all organs to different degrees (Supplemental Material). Therefore, dietary effects in an organ depend on sex and the genetic context.

### 2.8 Correlations with blood lipids

We wondered whether the full blood (or blood plasma) itself, as an easily accessible sample, is a good proxy for lipidomic changes occurring in other tissues and organs. To investigate whether lipidomic changes in mouse organs are reflected in blood samples, we performed a correlation analysis of amounts of individual lipids in the different organs with the respective lipids in bodily fluids. We consider only significant (*p <* 0.05) positive correlations as indicative for corresponding changes in amounts. To identify which organs are best reflected by blood samples, these positive correlations were used as input for hierarchical clustering (Figure 6 A). Correlations in blood plasma are used as a reference point. The closer the organs cluster with blood plasma, the better their lipid composition is mirrored in the circulation. It appears that positive correlations of amounts of organ lipids with amounts of plasma lipids can be found across the entire lipidome and for all organs. Expectedly, full blood clusters closest with blood plasma since they are sharing the storage lipid complement (TAG and CE) organized in lipoprotein particles. Interestingly, it is liver that clusters closest with blood samples (blood plasma or full blood), followed by adipose tissue and muscle, while the brain and spleen lipidomes are most distant. Hence, changes in the liver lipidome are best reflected in blood samples, while lipidomic changes in brain are only weakly reflected by the blood lipidome. In liver, mostly changes in PC species correlate well with plasma samples. Additionally, changes in liver PI and TAG species are reflected in plasma samples. In muscle and adipose tissue, mostly TAG species are reflected in the plasma lipidome. It should be noted that organs were not subjected to perfusion after dissection. Therefore, any residual blood in these organs could affect correlations of their lipidomes with blood lipidomes. However, this does not seem to be the case, as lipidomes of organs well supplied with blood (such as spleen, intestine or lung) do not correlate well with blood lipidomes.

**Figure 6:**
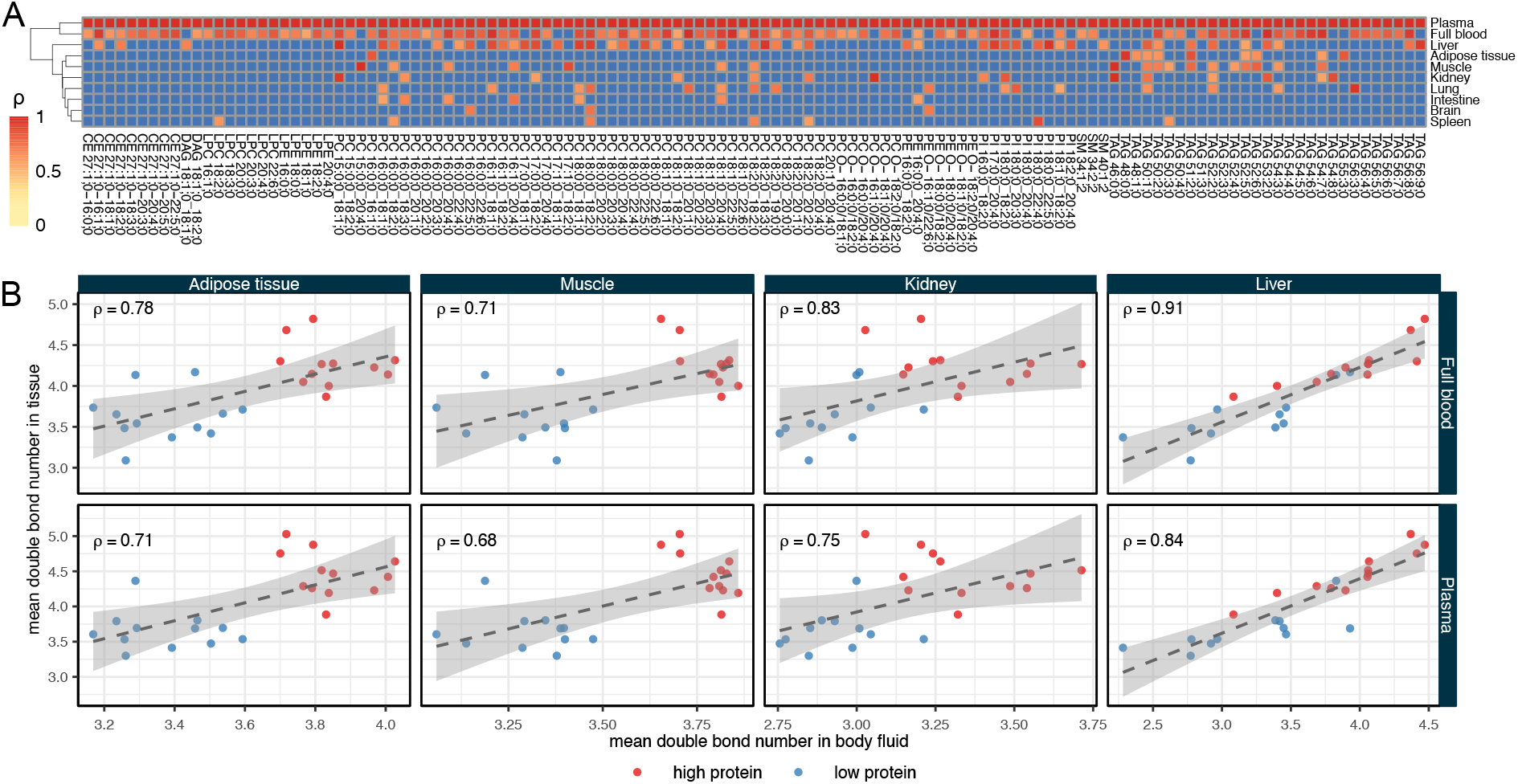
Correlations of organ lipidomes with blood lipidomes. A: Hierarchical clustering based on correlation coefficients for diet effects on individual lipids in different organs with lipids in blood plasma as reference point. Colour scale indicates correlation coefficient *ρ*. Non-significant positive correlations and negative correlations are shown in blue. B: Scatter plots showing correlations for TAG lipidomic features in organs with counterparts in blood plasma and full blood. Dashed lines show a linear regression of the data (with grey areas indicating the 95% confidence interval). In both panels only significant (*p <* 0.05) positive correlations are shown and correlations were adjusted for sex and genotype.

Considering correlations based on lipidomic features, TAG appears to be the class that is best reflected in the blood lipidome for various organs. For adipose tissue, muscle, liver, and kidney there are significant positive correlations of TAG unsaturation and chain length indices in both blood plasma and full blood (Figure 6 B). Additionally, there are robust correlations for PI features in liver, intestine, and kidney (not shown).

Even though there are numerous organ lipids whose abundance is reflected in blood samples, the blood lipidome is not a comprehensive mirror of lipidomic changes in organs. In most cases, not more than 5% of lipids in an organ are positively correlated with their amounts in the circulation. The exception is liver, with ca. 15% of lipids correlating with blood. Therefore, organ lipidomes respond in a specific manner to different stimuli and the responses are not necessarily reflected in blood.

## 3 Discussion

Lipidomics has become an indispensable technology in basic, clinical, and pharmaceutical research to investigate lipid metabolism quantitatively at the molecular level. Therefore, a thorough characterization of experimental model systems is required to provide a reference for future studies, facilitating experimental design and data interpretation. Here, we present a comprehensive and quantitative lipidomic atlas of mouse organs, full blood, and blood plasma. This study confirms a remarkable specificity of the organ’s lipid composition both at the level of lipid classes as well as for the individual lipid molecules. In fact, each organ shows a unique and unambiguous lipid fingerprint. Including mice of different genotypic background, sex and feeding of two different diets allowed for an analysis of lipidomic flexibility for each organ and lipid class. This analysis reveals that the lipidome of certain organs is more susceptible to variation of genotype, sex, and diet while other organs exhibit a remarkably robust lipid composition unimpeded by these perturbations. The multi-factorial study design further enabled the analysis of genotype- and sex-specific diet effects for each organ. Again, when specificity could be demonstrated, it was organ- and lipid class-dependent, highlighting the complexity of an individual’s lipid metabolism. Moreover, correlations of organ lipidomes with full blood and plasma lipidomes revealed that lipidomic changes in the brain, spleen, and intestine are only poorly mirrored in the circulation, while the liver lipidome is well reflected in the blood.

Most importantly, the factorial design of this study enabled the analysis of lipidomic plasticity of the different lipid classes in different organs (Figure 3) and the lipidomic flexibility of different organs in response to diet, sex, and genotype (Figure 4). We define lipidomic plasticity as the degree of structural heterogeneity/variation within a given lipid class, i.e., the possible range of the degree of unsaturation (double bond numbers) and hydrocarbon chain length. Lipidomic flexibility of an organ, on the other hand, is the magnitude of lipidomic changes induced by experimental conditions. Both parameters are characteristic features of lipidomes and specific for organs, tissues, and even cell types.

### 3.1 Organ-specific lipidomes

The specific lipid composition of organs has been documented in detail previously [Fitzner et al., 2020, Jain et al., 2014, Parker et al., 2019, Pradas et al., 2018, Symons et al., 2021]. The lipid composition obviously reflects histological and cellular structures, which are the foundation of the organ’s functions. For example, adipose tissue with its triglyceride- laden lipid droplets in their adipocytes is composed almost exclusively of triglycerides (ca. 99 mol%) [Grzybek et al., 2019]. Lung samples are characterized by high amounts of surfactant components, the most prominent being dipalmitoyl PC (PC 16:0/16:0), PG, and SM [Lang et al., 2005]. Brain samples are rich in PE O-, hexosyl ceramide, ceramide, and cholesterol. PE O-, hexosyl ceramide and cholesterol are major components of myelin sheaths, structures insulating the axons in neurons [Fitzner et al., 2020]. High concentrations of ceramides likely serve as precursor for the synthesis of complex glycolipids such as sulfatide and gangliosides, which are major lipid components of the mammalian brain [Schnaar et al., 2014]. These lipids were not included in our analysis. Furthermore, high amounts of CL in both liver and kidney reflect the large numbers of mitochondria in these organs.

Compositional specificity not only at the lipid class level, but also at the level of individual lipid molecules, is achieved by organ-specific expression of lipid biosynthetic genes. This is best exemplified for ceramide synthases, which show highly tissue-specific expression profiles In intestine, ceramide synthase 6 is expressed highest with a preference for myristic (C14:0) and palmitic (C16:0) acid, resulting in ceramides with a total carbon number of 32-34. In brain, ceramide synthase 1 is expressed at the highest levels with a preference for stearic acid (18:0), resulting in ceramides with a total carbon number of 36. At the other end of the scale is liver, in which ceramide synthase 2 is expressed abundantly [Laviad et al., 2008]. Ceramide synthase 2 prefers fatty acids with 20 to 26 carbons, giving rise to ceramide species with total carbon numbers ranging from 38 to 44. These expression patterns are reflected in the ceramide profiles for the various tissues (Figure 2). Similarly, organ-specific species profiles for other lipid classes are achieved by distinct expression and activity profiles of fatty acid desaturases, elongases and acyl transferases [Harayama et al., 2014]. However, there is much to be learned about the molecular mechanisms that give rise to the perplexing complexity of lipid metabolism and composition for different organs, tissues, and cell types.

### 3.2 Lipidome plasticity and flexibility and its implications for experimental design and data interpretation

The largest variation in the data set is caused by organ-specific differences of the lipidomes. Diet, genotype, and sex each contribute to about 5% of overall variance. Nevertheless, for each organ a significant fraction of the lipidome is affected by the tested conditions. However, some organs are more responsive than others. For example, the brain lipidome is only affected by genotype while diet and sex hardly have any significant effect. Therefore, the brain appears to have a very robust lipid composition. This is confirmed by the very low internal plasticity of the major brain lipid classes (like for example PC, PE O-, hexosyl ceramide, Figure 3). Other sample types such as liver and full blood are affected by all the tested conditions. One may interpret a low lipidomic flexibility as a consequence of a tight coupling of composition, structure, and function. A very specific lipid composition (i.e., a low lipidomic plasticity) with particular structural features such as degree of unsaturation and fatty acid chain length are required to fulfill a specific function. In brain that would be the formation of the myelin sheath [Yurlova et al., 2011] and in the lung the coating of the alveolar interior with surfactant lipids like dipalmitoyl PC [Postle et al., 2001]. Liver and full blood on the other hand fulfill rather metabolic than structural functions, which requires/results in a greater flexibility to serve as buffer against metabolic perturbations.

Therefore, we propose lipidomic plasticity and flexibility as intrinsic properties of an organ or bodily fluid. Future studies will likely necessitate the extension of this concept to tissues and even cell types. Lipidomic plasticity and flexibility of an organ have important implications for experimental design: If an organ with high flexibility (resulting in large effect sizes) is under investigation, lower numbers of biological replicates might be sufficient to observe robust phenotypes. Furthermore, changes observed in organs that are characterized by low plasticity and/or flexibility might be more meaningful and hint at a severe phenotype. Another level of complexity is added by the fact that intervention effects (here simulated by diet) might depend on genotype or sex of the animal. This might be generalized to other backgrounds, like for example diet- or genotype-specific drug effects in pharmaceutical time-dose studies.

Blood samples (in most cases blood plasma) are usually used to study intervention effects in model systems. However, and perhaps not surprisingly, our results suggest that not every organ’s lipid metabolism is reflected in the circulation. As the liver plays a central role in lipoprotein metabolism, the liver lipidome correlates strongest with blood plasma, confirming previous studies [Parker et al., 2019, Sprenger et al., 2021]. Lipid metabolism of other organs is not well reflected in blood, especially brain, spleen, and intestine. However, more research is obviously needed to understand how the blood and plasma lipidome reflects systemic effects on lipid metabolism. Nevertheless, if organ-specific effects of interventions are eventually desired or expected, it is recommended to not only rely on an initial screening of blood samples as important phenotypes might be missed.

### 3.3 Limitations of the study

This study presents a complex analysis of mouse organ lipidomes and their dependence on different experimental conditions/factors. However, some important organs or bodily fluids were not included, such as reproductive organs, pancreas, heart muscle, brown adipose tissue, cerebrospinal fluid, or urine. Analysis of additional tissue and cell types from various organs will certainly increase the degree of complexity. Furthermore, the analysis of complex glycolipids would be insightful, as would the inclusion of additional mouse genotypes. Finally, the dietary intervention used here is rather subtle. Treatment with a high fat diet or different drugs might lead to more pronounced effects and hence different conclusions regarding organ flexibility and dependence on genotype and sex.

### 3.4 Conclusion

Here we present a systematic analysis of the lipid composition of mouse organs and blood samples and an assessment of overall plasticity and flexibility of the mouse lipidome induced by diet, genotype, and sex. This dataset provides a valuable resource for basic and pharmaceutical researchers using mouse as a model system and complements existing proteomic and transcriptomic datasets. More basic and quantitative characterizations of model systems are required for other omics technologies as they will aid standardization of research areas and inform experimental design and facilitate interpretation of lipidomic datasets.

## 4 Acknowledgements

The authors thank Melanie Weber and Steffi Lenhard for their excellent technical assistance and Marta Pasikowska from Przychodnia Weterynaryjna VIVA, Wrocław, Poland for providing blood collection tubes and advice on animal samples handling. CK gratefully acknowledges stimulating input from Serj Tankian.

## 5 Author contributions

MAS, KS, and CK conceived and supervised the project and designed experiments. MAS performed the experiments and acquired data. MJG and RH provided data processing and analysis tools and workflows. JH provided mouse strains. CK analyzed data and wrote the manuscript.

## 6 Supplemental Material

Supplemental table and figure captions are provided in the respective files.

**Supplemental Text** Supplemental text and figures for assessment of analytical method performance.

**Supplemental Table 1** Complete lipidome data, including matching of lipid names to SwissLipids identifiers.

**Supplemental Table 2** Data on mouse diet composition as provided by the manufacturer (Envigo).

**Supplemental Table 3** Results of linear regression analysis.

## 7 Materials & Methods

### 7.1 Mouse handling

Two standard laboratory mouse wild-type strains were selected: Hsd:ICR (CD-1^®^) and C57BL/6JOlaHsd, genetically representing an outbred and inbred population, respectively. The mice were allowed to breed within the strains, and once pregnancy was confirmed, the pregnant females were put on two standard, healthy diets differing by protein content: Teklad global 14% protein (Envigo #2014S, low protein) and Teklad global 18% protein (Envigo #2018S, high protein). The high protein diet delivered 24% of total calories from proteins and 18% from fats, whereas the low protein diet delivered 20% of total calories from proteins and 13% from fats. Qualitatively, the lipid compositions of high and low protein diets are comparable (Supplemental Material). After birth, mothers were kept at the same diet and the litter mice were introduced to the same diet as their mothers after weaning.

At the age of ca. 8.5 weeks, mice were weighted and then sacrificed, to yield three females and three males from each strain and each diet. Hsd:ICR (outbred) animals came from different litters. On the contrary, C57BL/6JOlaHsd (inbred) animals came from the same litter, that is all six mice (three males and three females) for each diet were siblings. All animals were maintained in pathogen-free conditions in the animal facility of Max Planck Institute of Molecular Cell Biology and Genetics (Dresden, Germany) and all handling was performed in accordance with German animal welfare legislation.

### 7.2 Sample preparation

Mice were sacrificed and immediately decapitated, and the full blood (F) samples were collected directly from necks into EDTA-tubes (Sarstedt, #41.1395.105). For plasma (P) preparation, tubes were centrifuged for 10 minutes at 2000 g and blood plasma was collected within 1 hour after blood collection. After diluting full blood and plasma 50*×* with water, samples were frozen at *−*80°C until lipid extraction.

Organs and tissues were dissected from fresh cadavers after all blood was drained (no perfusion was performed), resulting in the following samples: liver-H (a section of the middle part of the left lateral lobe); skeletal muscle-M (left soleus); brain-B (section of left hemisphere); kidney-K (whole left kidney, without fat tissue); adipose tissue-A (from the abdominal region); small intestine-I (whole duodenum); lung-L (section of middle region of left lung) and spleen-S (whole spleen). These organs and tissues were put into 2 mL microcentrifuge tubes, immediately frozen on dry ice, weighted and stored at *−*80°C until homogenization.

Homogenization was performed after samples were thawed on ice and resuspended in 1.5 mL of 4°C cold 150 mM ammonium bicarbonate buffer (or with this buffer mixed 1:1 with ethanol, for adipose tissue samples [Grzybek et al., 2019]), by shaking in 2 ml tubes for 15 min (3 *×*5 minutes with 2 minutes cool down periods on ice) at 4°C with several 3.1 mm stainless steel beads using a Qiagen TissueLyser II.

Homogenized samples were diluted, and the following amounts were used for lipid extraction: 1 *µ*l blood plasma, 1 *µ*l full blood, 80 *µ*g brain, 100 *µ*g liver, 270 *µ*g intestine, 200 *µ*g kidney, 250 *µ*g lung, 650 *µ*g muscle, 400 *µ*g spleen, 100 *µ*g adipose tissue.

### 7.3 Analytical process design

Samples were divided into four separate analytical batches: plasma, adipose tissue and two batches for the remaining samples. Each batch was accompanied by a set of blank samples (pure buffer) and identical reference samples: human plasma for the plasma batch, full blood for the remaining sample types. These control samples were distributed evenly across each batch, extracted, and processed together with study samples to control for background signals and intra-run reproducibility. All batches were measured on 4 consecutive days.

### 7.4 Lipid extraction

Lipid extraction for lipidomic analysis was performed as described for plasma [Surma et al., 2015] and adipose tissue [Grzybek et al., 2019]. For all other types of samples, the procedure was as follows. Lipids were extracted using a two-step chloroform/methanol procedure [Ejsing et al., 2009] with chloroform:methanol 10:1 (V:V) and 2:1 (V:V) in the first and second step, respectively [Sampaio et al., 2011]. Prior to extraction, samples were spiked with an internal lipid standard mixture containing: cardiolipin 16:1/15:0/15:0/15:0 (CL, 50 pmol per extraction), ceramide 18:1;2/17:0 (Cer, 30 pmol), diacylglycerol 17:0/17:0 (DAG, 100 pmol), hexosyl ceramide 18:1;2/12:0 (HexCer, 30 pmol), lyso-phosphatidate 17:0 (LPA, 30 pmol), lyso-phosphatidylcholine 12:0 (LPC, 50 pmol), lyso-phosphatidylethanolamine 17:1 (LPE, 30 pmol), lyso-phosphatidylglycerol 17:1 (LPG, 30 pmol), lyso-phosphatidylinositol 17:1 (LPI, 20 pmol), lyso-phosphatidylserine 17:1 (LPS, 30 pmol), phosphatidate 17:0/17:0 (PA, 50 pmol), phosphatidylcholine 17:0/17:0 (PC, 150 pmol), phosphatidylethanolamine 17:0/17:0 (PE, 75 pmol), phosphatidylglycerol 17:0/17:0 (PG, 50 pmol), phosphatidylinositol 16:0/16:0 (PI, 50 pmol), phosphatidylserine 17:0/17:0 (PS, 100 pmol), cholesterol ester 20:0 (CE, 100 pmol), sphingomyelin 18:1;2/12:0;0 (SM, 50 pmol), triacylglycerol 17:0/17:0/17:0 (TAG, 75 pmol) and cholesterol D6 (Chol, 300 pmol). After extraction, the organic phase was transferred to an infusion plate and dried in a speed vacuum concentrator. 1st step dry extract was re-suspended in mM ammonium acetate in chloroform/methanol/propanol (1:2:4, V:V:V) and 2nd step dry extract in 33% ethanol solution of methylamine in chloroform/methanol (0.003:5:1; V:V:V). All liquid handling steps were performed using Hamilton Robotics STARlet robotic platform with the Anti Droplet Control feature for organic solvents pipetting. All solvents and chemicals used were of analytical grade.

### 7.5 Mass spectrometry and data processing

Mass spectrometry analysis was performed as described previously for plasma [Surma et al., 2015] and adipose tissue [Grzybek et al., 2019]. Samples were analyzed by direct infusion on a QExactive mass spectrometer (Thermo Scientific) equipped with a TriVersa NanoMate ion source (Advion Biosciences). Samples were analyzed in both positive and negative ion modes with a resolution of *R*_*m/z*=200_ = 280000 for MS and *R*_*m/z*=200_ = 17500 for MSMS experiments, in a single acquisition. MSMS was triggered by an inclusion list encompassing corresponding MS mass ranges scanned in 1 Da increments [Surma et al., 2015]. Both MS and MSMS data were combined to monitor CE, DAG and TAG ions as ammonium adducts; PC, PC O-, as acetate adducts; and CL, PA, PE, PE O-, PG, PI and PS as deprotonated anions. MS only was used to monitor LPA, LPE, LPE O-, LPI and LPS as deprotonated anions; Cer, HexCer, SM, LPC and LPC O-as acetate adducts and cholesterol as ammonium adduct of an acetylated derivative [Liebisch et al., 2006]. Data were analyzed with in-house developed lipid identification software based on LipidXplorer [Herzog et al., 2011, 2012].

Data post-processing and normalization were performed using an in-house developed data management system. Only lipid identifications with a signal-to-noise ratio *>* 5, and a signal intensity 5-fold higher than in corresponding blank samples were considered for further data analysis.

### 7.6 Lipid nomenclature

Lipid molecules are identified as species or subspecies. Fragmentation of the lipid molecules in MSMS mode delivers subspecies information, i.e., the exact acyl chain (e.g., fatty acid) composition of the lipid molecule. MS only mode, acquiring data without fragmentation, cannot deliver this information and provides species information only. In that case, the sum of the carbon atoms and double bonds in the hydrocarbon moieties is provided. Lipid species are annotated according to their molecular composition as lipid class <sum of carbon atoms>:< sum of double bonds>;< sum of hydroxyl groups>. For example, PI 34:1;0 denotes phosphatidylinositol with a total length of its fatty acids equal to 34 carbon atoms, total number of double bonds in its fatty acids equal to 1 and 0 hydroxylations. In case of sphingolipids, SM 34:1;2 denotes a sphingomyelin species with a total of 34 carbon atoms, 1 double bond, and 2 hydroxyl groups in the ceramide backbone. Lipid subspecies annotation contains additional information on the exact identity of their acyl moieties and their *sn*-position (if available). For example, PI 18:1;0_16:0;0 denotes phosphatidylinositol with octadecenoic (18:1;0) and hexadecanoic (16:0;0) fatty acids, for which the exact position (*sn*-1 or *sn*-2) in relation to the glycerol backbone cannot be discriminated (underline “_” separating the acyl chains). On contrary, PC O-18:1;0/16:0;0 denotes an ether-phosphatidylcholine, in which an alkyl chain with 18 carbon atoms and 1 double bond (O-18:1;0) is ether-bound to *sn*-1 position of the glycerol and a hexadecanoic acid (16:0;0) is connect via an ester bond to the *sn*-2 position of the glycerol (slash “/” separating the chains signifies that the *sn*-position on the glycerol can be resolved). Lipid identifiers of the SwissLipids database [Aimo et al., 2015] are provided in Supplemental Material.

### 7.7 Data analysis

Data were analyzed with R version 4.0.3 [R Core Team, 2020] using tidyverse (v.1.2.1) packages [Wickham, 2020]. Data were corrected for batch effects and analytical drift based on reference samples. Only lipids with amounts *>* 1 pmol and identified in at least two out of three biological replicates were used for further data analysis, resulting in 796 lipid (sub-)species included in the final dataset. Molar amount values (in pmol) of individual lipid (sub-)species were normalized to total lipid content per sample, yielding molar fraction values, expressed in mol%.

A weighted mean length per total fatty acids 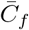 or weighted mean saturation per total fatty acids 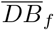 per lipid classes is calculated [Tam et al., 2021]. This returns an estimate of the mean total fatty acid length or mean total number of fatty acid double bonds in this lipid class:

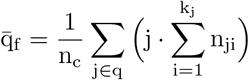

where

- 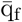 is the weighted mean length/saturation per hydrocarbon moiety,
- q = {q_1_, …, q_max_} is the feature of a lipid with a list of instances: e.g. total double bonds = {0, 1, 2, …, 8} or total length = {34, 35, 36, …, 42},
- *n*_*c*_ is the total molar amount of the respective lipid class,
- *n*_*ji*_ is the molar amount of lipid species *i* with the respective matching the feature instance *j* and
- *k*_*j*_ is the number of species within feature instance *j*.

Principal component analysis (PCA) was performed using the stats::prcomp() function on centered and scaled data.

The internal lipid class plasticity was calculated based on the minimum-maximum range of values for weighted mean of lipid class unsaturation and the weighted mean of lipid class carbon chain length. The ranges for each feature across different sample types was scaled to assume values between 1 and 10, meaning that the class with the widest range across organs receives a value of 10, the class with narrowest range receives a value of 1. Features with intermediate ranges receive values between 1 and 10, according to scale. The scaled ranges were then used to calculate the internal lipid class plasticity by multiplying the scaled carbon chain length range with the scaled double bond number range (Figure 3). Because of the limited lipid composition, adipose tissue was omitted from the plasticity calculations.

Sex, diet and genotype effect sizes (*β* coefficients) were determined for centered and scaled lipidomic features (z scores for lipid class amount in mol%, weighted mean of lipid class unsaturation and weighted mean of lipid class carbon chain length) by linear regression using the stats::lm() function adjusting for the other covariates. For the calculation of sex- and genotype-specific diet effects, the respective interaction terms were included in the linear regression. Mouse plots were created with the R library gganatogram (v.2) [Maag, 2018].

Diet-dependent Spearman correlation coefficients (*ρ*) between blood and tissue samples were calculated using the function RVAideMemoire::pcor.test (v.0.9-78) adjusting for sex and genotype [Hervé, 2020]. Hierarchical clustering (Euclidian distance, complete linkage) heatmap generation was performed using the pheatmap::pheatmap() (v.1.0.12) function, which was also used for heatmap generation [Kolde, 2019].

## Footnotes

1 Lipid class abbreviations used throughout the text: TAG, triglycerides; CL, cardiolipin; DAG, diacylglycerol; LPA, lysophosphatidate; LPC, lyso-phosphatidylcholine; LPC O-, lyso-phosphatidylcholine ether; LPE, lyso-phosphatidylethanolamine; LPE O-, lyso-phosphatidylethanolamine ether; LPG, lyso-phosphatidylglycerol; LPI, lyso-phosphatidylinositol; LPS, lyso-phosphatidylserine; PA, phosphatidate; PC, phosphatidylcholine; PC O-, phosphatidylcholine ether; PE phosphatidylethanolamine; PE O-, phosphatidylethanolamine ether; PG, phosphatidylglycerol; PI, phosphatidylinositol; PS, phosphatidylserine; CE, cholesterol ester; SM, sphingomyelin; ST, cholesterol; HexCer, hexosyl ceramide; Cer, ceramide.

