## Supplemental Text for "Flexibility of a mammalian lipidome — insights from mouse lipidomics"

### Flexibility of a mammalian lipidome – insights from mouse lipidomics: Supplemental material

#### Assessment of method performance

Method performance of the shotgun lipidomics technology for mouse tissues was assessed for brain and liver samples. Dynamic range, limit of detection (LOD) and limit of quantification (LOQ) are lipid class-specific parameters, which depend on differential extraction, ionization, and fragmentation efficiencies of the various lipid classes. In order to determine these parameters, lipid class-specific standards were titrated to a fixed amount of tissue homogenate. Linearity and proportionality were assessed for 20 lipid classes by linear regression of log-transformed lipid amounts and their intensities, and reported as $R^2$ and slope, respectively. For liver samples, both, linearity, and proportionality were excellent across a range of up to 4 orders of magnitude for all lipid classes, with values close to 1 for $R^2$ and slope, respectively (Table 1). LOD and LOQ were determined by weighted linear regression (with weights being $1/x^2$) based on a signal-to-noise ratio of 3 for LOD and 10 for LOQ. For most lipid classes, LOD is around 1 pmol (Table 1). Provided that an optimal sample amount of mouse tissue yields ~10,000 pmol of total lipid, 1 pmol of a given lipid molecule is theoretically detectable in a typical sample, which corresponds to 0.01 mol%. Similar results were obtained for brain samples (not shown).

The reproducibility of the shotgun lipidomics method for tissue samples was assessed by 6 repeated, independent measurements of identical aliquots of liver and brain samples. For the assessment of reproducibility, only lipids present in at least 4 out of 6 replicates were considered. For liver samples, 327 lipids fulfilled this requirement. The relative standard deviation (RSD) of each lipid molecule was calculated and plotted against its abundance (in pmol, Figure 1) and the data revealed an inverse correlation between lipid abundance and RSD. The median technical variation was 5.6%, while 84.7% of the detected lipids showed an RSD <15%. For brain samples, the median technical variation was 6.3%, while 84.6% of the detected lipids showed an RSD <15% (not shown).

*Table 1: Limit of detection (LOD) and limit of quantification (LOQ) of various lipid classes in liver tissue.*

| class | LOD (pmol) | LOQ (pmol) | slope | $R^2$ |
|---|---|---|---|---|
| Cer | 1.0 | 3.1 | 0.88 | 0.99 |
| ST | 1.2 | 3.5 | 0.95 | 0.98 |
| CL | 1.3 | 3.9 | 0.89 | 0.99 |
| HexCer | 0.8 | 2.3 | 0.93 | 0.99 |
| LPA | 0.6 | 1.9 | 1.05 | 0.99 |
| LPC | 0.6 | 1.9 | 0.97 | 0.99 |
| LPE | 0.8 | 2.3 | 0.98 | 0.99 |
| LPG | 0.6 | 1.8 | 0.93 | 1.00 |
| LPI | 0.7 | 2.1 | 1.07 | 0.98 |
| LPS | 0.8 | 2.5 | 1.04 | 0.99 |
| SM | 0.8 | 2.5 | 0.94 | 0.99 |
| TAG | 0.6 | 1.8 | 0.95 | 1.00 |
| CE | 0.7 | 2.0 | 0.92 | 0.99 |
| DAG | 0.6 | 1.9 | 0.95 | 1.00 |
| PA | 0.6 | 1.9 | 1.09 | 0.99 |
| PC | 1.5 | 4.7 | 0.84 | 0.97 |
| PE | 1.2 | 3.6 | 0.89 | 0.99 |
| PG | 1.1 | 3.2 | 0.86 | 0.99 |
| PI | 0.6 | 2.0 | 1.02 | 0.99 |
| PS | 1.1 | 3.5 | 1.32 | 0.96 |

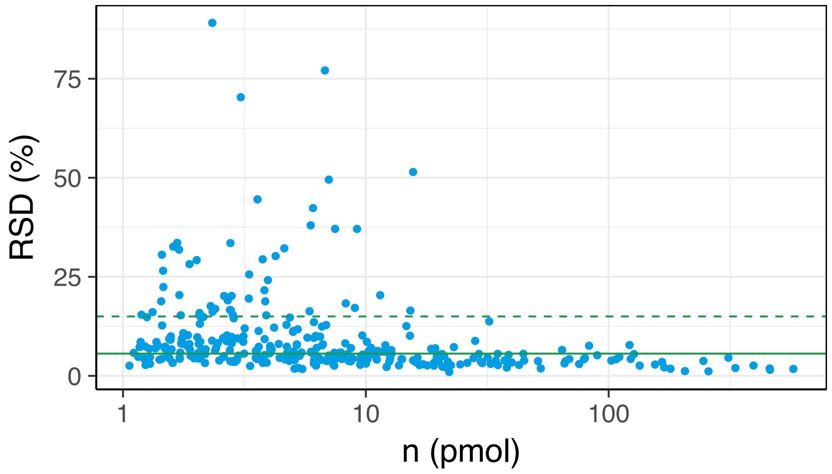


*Figure 1: Relative standard deviation (RSD) calculated for 327 lipids present in at least 4 out of 6 replicates of mouse liver sample. Each lipid molecule RSD was calculated and plotted against its abundance (in pmol). The median technical variation is 5.6% (green solid line) and RSD <15% (green dotted line) was observed for 84.7% of the detected lipids.*
